# First report of antiviral activity of nordihydroguaiaretic acid against Fort Sherman-like virus (Orthobunyavirus)

**DOI:** 10.1101/2020.08.04.236133

**Authors:** Florencia Martinez, María Laura Mugas, Juan Javier Aguilar, Juliana Marioni, Marta Silvia Contigiani, Susana C. Núñez Montoya, Brenda S. Konigheim

**Affiliations:** Universidad Nacional de Córdoba, Facultad de Ciencias Médicas, Instituto de Virología “Dr. J. M. Vanella”-, Argentina. Enfermera Gordillo S/N, Ciudad Universitaria, X5000HUA Córdoba, Argentina; Consejo Nacional de Investigaciones Científicas y Técnicas (CONICET), Argentina; Universidad Nacional Córdoba, Facultad Ciencias Químicas, Dpto. Ciencias Farmacéuticas, Argentina. Haya de la Torre y Medina Allende, Ciudad Universitaria. X5000HUA Córdoba, Argentina; CONICET, Instituto Multidisciplinario de Biología Vegetal (IMBIV). Av. Vélez Sarsfield 1666. CP: X5016GCN Córdoba, Argentina

**Author notes:** Corresponding Author: Correspondence/Mail should be addressed to Brenda S. Konigheim, Universidad Nacional de Córdoba, Facultad de Ciencias Médicas, Instituto de Virología “Dr. J. M. Vanella”, Argentina. Enfermera Gordillo S/N, Ciudad Universitaria, X5000HUA Córdoba, Argentina., Consejo Nacional de Investigaciones Científicas y Técnicas (CONICET), Argentina., / /.

**Keywords:** nordihydroguaiaretic acid, Fort Sherman virus, antiviral activity, lipid metabolism

## Abstract

The genus Orthobunyavirus are a group of viruses within arbovirus, with a zoonotic cycle, some of which could lead to human infection. A characteristic of these viruses is their lack of antiviral treatment or vaccine for its prevention. The objective of this work was to study the *in vitro* antiviral activity of nordihydroguaiaretic acid (NDGA), the most important active compound of *Larrea divaricata* Cav. (*Zigophyllaceae*), against Fort Sherman-like virus (FSV-like) as a model of Orthobunyavirus genus. At the same time, the effect of NDGA as a lipolytic agent on the cell cycle of this viral model was assessed. The method of reducing plaque forming units on LLC-MK2 cells was used to detect the action of NDGA on CbaAr426 and SFCrEq231 isolates of FSV-like. NDGA did not show virucidal effect, but it had antiviral activity with a similar inhibition in both isolates, which was dose dependent. It was established that the NDGA has a better inhibition one-hour post infection (p.i.), showing a different behavior in each isolate, which was dependent upon the time p.i. Since virus multiplication is dependent on host cell lipid metabolism, the antiviral effect of NDGA has been previously related to its ability to disturb the lipid metabolism, probably by interfering with the sterol regulatory element-binding proteins (SREBP) pathway and the 5-lipoxigenase (5-LOX). We determined by using caffeic acid, a 5-LOX inhibitor, that the inhibition of this enzyme negatively affected the FSV-like replication; and by the use of resveratrol, a SREBP1 inhibitor, it was showed that the negative regulation of this pathway only had action on the SFCrEq231 reduction. In addition, it was proved that the NDGA acts intracellularly, since it showed the ability to incorporate into LLC-MK2 cells. The information provided in this work converts the NDGA in a good antiviral candidate, especially for Orthobunyavirus infections, and a useful tool for the biochemical study of FSV-like that causes an infection poorly studied and potentially dangerous.

## 1. Introduction

The Orthobunyavirus genus (Peribunyaviridae) includes 88 species, some of which are important pathogens not only for humans but also for animals, and they are maintained in nature by zoonotic cycles. The main prototype member of this genus is the Bunyamwera virus which is associated with infections in human (feverish disease that occasional involvement nervous system) and animals (miscarriage and teratogenic effects) (Strauss and Strauss, 2008, Rodrigues Hoffmann et al., 2013). Another relevant specie is the Cache Valley virus, endemic to North America, and which causes profoundly serious diseases in ruminants with consequent economic losses (Waddell et al., 2019). Fort Sherman virus (FSV), other species of this genus, was isolated in 1985 from a US soldier with acute febrile disease based in Panama (de Oliveira Filho et al., 2020). In Argentina, has been registered circulation of viruses of this genus associated with febrile disease in humans (Tauro et al., 2012) and isolation has been obtained from dead equines with neurological signs and from the brain of an equine abortion (Tauro et al., 2015). This isolates were originally classified as Cache valey, but they are now considered different variants of Fort Sherman virus (FSV-like) since only the amino acid sequence of these isolates is known and not the complete genome (de Oliveira Filho et al., 2020). Thus, Orthobunyavirus expand their distribution area as a potential threat to animal and human health.

Members of the Orthobunyavirus are characterized by having a single-stranded negative-sense genome RNA divided into three segments (L, M and S), wrapped in a lipid bilayer. The segment S-mRNA encodes protein N and non-structural protein NSs, with overlapping reading frames (Elliott, 2014). M-mRNA segment encodes a polyprotein that is post-translationally cleaved into the nonstructural protein NSm and Gn and Gc glycoproteins (Gentsch and Bishop, 1979). L-mRNA encodes RNA-dependent RNA polymerase (Elliott, 2014). These common genetic traits in the genus are especially useful when using any of these viruses (species) as a model to evaluate potential antiviral agents against any Orthobunyavirus.

Since there are no vaccines or antiviral therapies available to treat Orthobunyavirus infections, the development of new antiviral strategies is urgently required (Hover et al., 2016). Medicinal and aromatic plants have been used as medicines and, with the development of science, represent a source of chemical diversity, which have provided not only important therapeutic compounds but also drugs protype (Tagboto and Townson, 2001; Clardy and Walsh, 2004). In this regard, the great diversity of Argentine plants, offers interesting possibilities for finding novel antiviral compounds (Visintini et al., 2013). A South American species extensively studied because of its multiple uses in folk medicine is *Larrea divaricata* Cav. (*Zigophyllaceae*), popularly known as “jarilla”. The most important active substance of this vegetal species is the nordihydroguaiaretic acid (NDGA), which is an exclusive secondary metabolite of this genus (Hyder et al., 2002).

The antiviral effect of NDGA and its derivatives has been reported in numerous studies, and the range of evaluated viruses is extensive (Gnabre et al., 1995; Huang et al., 2003; Hwu et al., 2008; Uchide et al., 2005; Park et al., 2003; Craigo et al., 2000; Konigheim et al., 2012). Some authors have associated the antiviral effect of NDGA with an inhibition of lipogenesis in some viruses, including arboviruses, which is necessary during viral replication. Specifically, it inhibits the protein binding to sterile regulatory elements (SREBP), which regulates lipid biosynthesis and homeostasis in mammals (Merino-Ramos et al., 2017; Syed and Siddiqui, 2011). NDGA also inhibits the lipoxygenase (5-LOX), which converts arachidonic acid into oxygenated products with important biological properties (Bhattacherjee et al., 1988). Consequentially, the background leads us to think that NDGA could be a possible compound with broad-spectrum antiviral activity and it would be involved in the inhibition of the biosynthesis of lipids, that could by necessary in the replication of the Orthobunyavirus.

In this work, the objective was to expand the knowledges on the antiviral activity of nordihydroguaiaretic acid on other arboviruses not previously tested. Two isolates of the FSV-like were used as a model of Orthobunyavirus genus, with the hypothesis that the virus needs the cellular lipid machinery for its replication and, therefore, NDGA is a possible antiviral candidate.

## 2. Material and methods

### 2.1. Compounds and standard solutions

The nordihydroguaiaretic acid (NDGA) was obtained from *Larrea divaricata* Cav. (*Zigophyllaceae*) with purity of 95 % determined by HPLC-UV-Vis (supplemental Fig. 1). Standard solution of NDGA was 33 mM. Ribavirin (Filaxis, Argentina) was used as positive control (C+) for RNA virus-antiviral activity (Diamond et al., 2002; Livonesi et al., 2006; Takhampunya et al., 2006) and was used at the same concentration which the NDGA was evaluated (90 µM) that is not cytotoxic (supplemental Fig. 2). Resveratrol (Sigma-Aldrich) 8 mM. The stock solution of caffeic acid (Sigma-Aldrich) was prepared in a concentration of 80 mM. All the standar solutions were prepared in dimethylsulfoxide (DMSO, Tetrahedron, USA). The concentration of DMSO in the assays was not greater than 1 %.

### 2.2. Cell cultures and viruses

*Macaca mulatta* monkey kidney cells (LLC-MK2 ATCC^®^ CCL-7) was used as host. They were grown in humidity atmosphere at 37 °C with 5% CO_2_. Eagle’s minimal essential medium (EMEM, Gibco, USA) supplemented with 10% (v/v) Fetal Bovine Serum (FBS, Natocor, Argentina), L-glutamine (30 μg/mL, Sigma-Aldrich, USA) and gentamicin (50 μg/mL, Sigma-Aldrich), was used as a growth medium (GM). The same formulation with 2% FBS was used as a maintenance medium (MM).

The following isolates of Fort Sherman-like virus (FSV-like) were used: CbaAr426 and SFCrEq231 (Bianchini et al., 1968; Tauro et al., 2015). Each viral stock was obtained and titrated according to the methods described by Contigiani and Sabattini (1977). The viral titration was expressed in plaque forming unit per milliliter (PFU/mL), being 1.2 × 10^6^ PFU/mL for CbaAr426 and 1.2 × 10^7^ PFU/mL for SFCrEq231.

### 2.3. *In vitro* cytotoxicity assay

In order to use non-toxic concentrations for the host cells when the antiviral activity is assessed, previously, cytotoxicity tests were carried out on these cells. By means of the Neutral Red (NR) uptake assay the cellular viability (CV) was measured (Neutral Red, Gibco, USA), following a procedure previously described by other authors (Borenfreund and Puerner, 1985; Ooi et al., 2004). Different dilutions of NDGA were added to a confluent monolayer of LLC-MK2 cells, using three replicates of each concentration, which were incubated at 37 °C in 5% CO_2_ for 72 h. Cells incubated only with MM (containing < 1% DMSO) were used as cell controls (CC, n=3). The absorbance of the extracted NR from cellular inside was measured at 540 nm by using a microplate reader (BioTek ELx800, USA). The percentage of CV (% CV) was calculated with respect to CC (100 % CV). By using the Origin 8.6 software, the curves % CV *vs* concentrations of NDGA were plotted with a non-linear regression analysis (Sigmoidal curve, R^2^ > 0.9).

From these graphs, the following parameters of toxicity were estimated: the concentration that reduced the viable cells to 50% (CC_50_), a subtoxic concentration (SubTC) specified as the concentration that causes 20% cellular death and produces slight morphologic changes observed by microscopy (Cholewa et al., 1994), and the Maximum Non-Cytotoxic Concentration (MNCC) defined as the maximum concentration of sample that exhibits more than 90% viable cells and exerts no cytotoxic effect detected by microscopic monitoring (Liu, et al., 2009).

### 2.4. Plaque reduction assay

This methodology was used to quantify viable virus in all trials described below, counting the lysis plaques produced by a viral particle in the monolayer cells used as host. Thus, the ability of the FSV-like to produce lysis plaques in LLC-MK2 cells was seize to quantify viral inhibition by perfoming the technique of plaque forming units (PFU/mL). This technique consists of covering the cell monolayer (infected or not, treated, or untreated) with a nutrient semi-solid medium, prepared with equal parts of 1% agarose (Invitrogen, USA) and MM of double concentration. The scientific rationale behind this trial is to consider that when a virus infects a cell, its progeny can only migrate to the immediately neighboring cells, and not to the remote ones, since the semi-solid medium limits its mobility. Thus, each plate is considered initiated by a simple viral particle, so that it constitutes a clone (Del Barrio Alonso and Parra, 2000). Then, the cell monolayer was fixed with 10% formalin (Anedra, Argentina) during 2 h. The overlay medium was removed and the formed PFUs were revealed with 1 % Crystal Violet (Anedra, Argentina), according to the methods described previously (Cheng et al., 2008).

#### 2.4.1. Viral inactivation activity (*in vitro* virucidal activity)

The CC_50_ of NDGA was mixed with viral stock in 1:1 proportion (T: Treatment). The same amount of viruses in MM without compound (Viral Control, VC) was used to compare with T, and a solution of NDGA (CC_50_ in MM) was included as the Drug Cytotoxicity Control (DCC). The T and the two controls were incubated at 37 °C in 5% CO_2_ for 1 h. Then, a serial dilution (factor 10) of the T and controls were performed; and each dilution was inoculated in duplicate on a confluent monolayer of LLC-MK2 cells (3.5 x 10^5^ cells/mL). After incubation at 37 °C in 5% CO_2_ for 1 h, the semi-solid medium was added, and the incubation was restarted to complete 4 days. The infectivity of the remaining viruses after the treatment was determined by the PFU reduction regarding the PFU developed in VC, by performing the plaque reduction assay (Cheng et al., 2008).

#### 2.4.2. *In vitro* antiviral activity

Viral suspension (1000 PFU/well approximately) was inoculated in triplicate on a confluent monolayer of LLC-MK2 cells (3.5 × 10^5^ cells/mL To produce the adsorption of viral particle, the 24-well microplate was incubated at 4 °C for 1 h and then, cells were washed with phosphate buffered saline (PBS). Subsequently, MM was added, and the microplate was incubated at 37 °C in 5% CO_2_ for 1 h; this process allowed the synchronized penetration of the viral particle. After that, cells were overlaid with semi-solid medium containing consecutive serial 1:2 dilutions from SubTC VC and DCC at the same dilution concentrations were also included in triplicate. The microplate was incubated for 4 days. Results were expressed as inhibition percentage (I%) in correlation with the VC (100 % of viable virus) (Gescher et al., 2011).

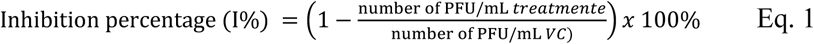

The I% is plotted as a function of the NDGA concentration, and the effective concentration that produces the 50 I% (EC_50_) was estimated by extrapolation.

#### 2.4.3. Kinetic curve of viral replication

To approximate the duration of the replication cycle of the FSV-like in the LLC-MK2 cells, the viral load was evaluated at different times after internalization of the viral particles. For that purpose, it was first necessary to guarantee a synchronized adsorption and internalization. Different cell monolayers were infected with each isolate of FSV-like separately (MOI ≈ 1) and incubated at 4 °C for 1 h to synchronize the infection. Three washes were then performed with PBS, to remove the virus that did not bind to the host cell. After adding MM to each well, an incubation was carried out (37 °C with 5 % CO_2_) for 66 h. At different times during this incubation (between 0 and 66h), labeled as post internalization (p.i.) times, the supernatants were collected and titled by means of the PFU assay. The objective of this test is to know the time that must elapse to detect a new viral progeny, so that the action of the NDGA can be evaluated at different times before the departure of the new virions.

#### 2.4.4. Evaluation of the NDGA effect on different times of infection

To study the effect of NDGA on different stages of FSV-like infection, the host cells were treated with NDGA before the cellular infection (pre-treatment, to assess whether the compound has an effect on cells that prevents, in any way, their infection), and at different times during the virus replication cycle (absorption, internalization, and post-internalization) (Sun et al., 2018).

The NDGA (MNCC) was added to a monolayer of LLC-MK2 cells (3.5 × 10^5^ cells/mL) at different times of infection with FSV-like (1000 PFU/mL aprox.): a) pre-treatment, the host cells were incubated with NDGA during 1 h at 37 °C before the infection; b) absorption, placing the compound together with the virus and incubating for 1 h at 4 °C; c) internalization, after the virus was adsorbed for 1 h at 4 °C, the infected monolayer is washed twice with PBS to eliminate the non-adsorbed viral particles, and the NDGA was placed and incubated for 1 h at 37 °C d) post-internalization, once the virus absorption (1 h at 4 °C, followed by two washes with PBS) and internalization (1 h at 37 °C in MM) were carried out, the NDGA was added with the semi-solid medium, and it was left to incubate (37 °C and 5 % CO_2_) until the end of the trial (control of antiviral activity already tested). In the experimental conditions a), b) and c), the NDGA only remained for 1 h and then, two sterile PBS washes were performed to eliminate the treatment. In the pre-treatment (a), the virus was added after these washes, allowing its adsorption and internalization. Finally, the semi-solid medium with out treatment was added to the three experimental conditions (a, b and c) for the final incubation at 37 °C with 5 % CO_2_ until the formation of PFU.

Different post-internalization (p.i.) times were also assessed to complete the study of the NDGA effect on the virus replication cycle of the two FSV-like isolates. The times were chosen according to the replication curves on the host cells, in order to evaluate those times prior to the release of the new viral progeny. Once the absorption and internalization of viruses were performed in LLC-MK2 cells, the NDGA (MNCC) was added at different times p.i. (1, 2, 3, 4, 5, 6, 7 and 8 h), contained in the semi-solid medium. And then, the multi well-plaque was left to incubate at 37 °C with 5 % CO_2_ for 4 days.

VC, DCC and C+ with ribavirin were included for each isntance. The viral titer was calculated by PFU/mL and the treatments at different times were compared, every hour, with the VC.

### 2.5. Evaluation of the lipogenesis inhibition in the infection by FSV-like

Resveratrol was used by its selective inhibition of the SREBP1, and caffeic acid for being a selective inhibitor of 5-LOX (Merino-Ramos et al., 2017; Koshihara et al., 1984). For this, both compounds were subjected to the methodology described above, with the aim to evaluate first their cytotoxicity in the host cells and then, their antiviral activity against the two isolates of FSV-like.

### 2.6. Statistical analysis

For the biological assays, the values were expressed in mean ± standard deviation (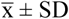) from three independent experiments, with three replicates inter trial. ANOVA and F-test (Origin 8.6 software) were used to assess the degree of statistical difference. Differences between means were considered significant at p < 0.05.

### 2.7 NDGA incorporation assay

The NDGA (993.3 µM, in excess to be able to detect it by HPLC) was added to a monolayer of LLC-MK2 cells (1.7 × 10^5^ cells/mL, growing for 48 h), and was incubated at 37 °C with 5 % CO_2_ for different times: 15, 30, 45, 60 min and 24 h in duplicate. At the end of each incubation time, the medium containing NDGA was harvested along with a washing of the cell monolayer with PBS (extracellular fraction, EC). Subsequently, the NDGA that entered into the cells was obtained at the same times (in duplicate), by the addition of CHCl_3_ (1 mL per well) to disrupt the cell membranes, and subsequent scraping to detach the cells (this procedure was repeated twice). Each well was stained with NR to ensure all cell monolayer was completely removed. Each sample obtained from each well was submitted to three freezing-thawing cycles, by using liquid nitrogen and a water bath at 40 °C, with the aim to break the cell structures and allow the release of the intracellular compound. Then, each sample was centrifuged for 15 min at 5000 rpm, once the pellet was separated, it was washed twice with CHCl_3_ and centrifuged again, and the three supernatants were pooled as the intracellular fraction (IC). CHCl_3_ was chosen because it not only disrupts cell membranes, but also helps to extract the NDGA due to its preferential solubility in this nonpolar solvent. So that, EC fractions were also partitioned with CHCl_3_ (1 mL) three times. All organic fractions (EC and IC) were evaporated to dryness in vacuum and dissolved in methanol grade HPLC (1 mL, MeOH-HPLC, Sintorgan) to obtain the corresponding solutions for subsequent analysis by HPLC (supplemental SA and SB).

## 3. Results

### 3.1. *In vitro* cytotoxicity assays

From the analysis of the plots % CV *vs.* concentration of NDGA in LLC-MK2 cells (supplemental Fig. 3), the values of the cytotoxic parameters were estimated as shown in Table 1.

**Table 1:** Values of cytotoxicity for Nordihydroguaiaretic acid in LLC-MK2 cells by means of the Neutral Red uptake assay.

| | Cytotoxic concentrations ($\mu\text{M}$) | | |
| --- | --- | --- | --- |
| | $\text{CC}_{50}^1$ | $\text{SubTC}^2$ | $\text{MNCC}^3$ |
| Nordihydroguaiaretic acid (NDGA) | $115.7 \pm 3.0$ | $99.0 \pm 4.9$ | $90.4 \pm 5.9$ |

Data are mean values from three independent experiments $\bar{x} \pm \text{SD}$ performed in triplicate.
<sup>1</sup> $\text{CC}_{50}$ : concentration that reduced the viable cells to 50 %.
<sup>2</sup>SubTC: subtoxic concentration, concentration that reduced the viable cells to 80 % ( $\text{CC}_{20}$ ).
<sup>3</sup>MNCC: Maximum Non-Cytotoxic Concentration.

### 3.2. Virucidal and antiviral activity

NDGA no showed viral inactivation (virucidal activity) in any of the isolates tested of the FSV-like. Nevertheless, a reduction in virus yield was observed when the *in vitro* antiviral activity of NDGA was assessed against two isolates of the FSV-like at different concentrations. The antiviral activity was dependent on the doses used in the treatment (Fig. 1).

**Figure 1:**
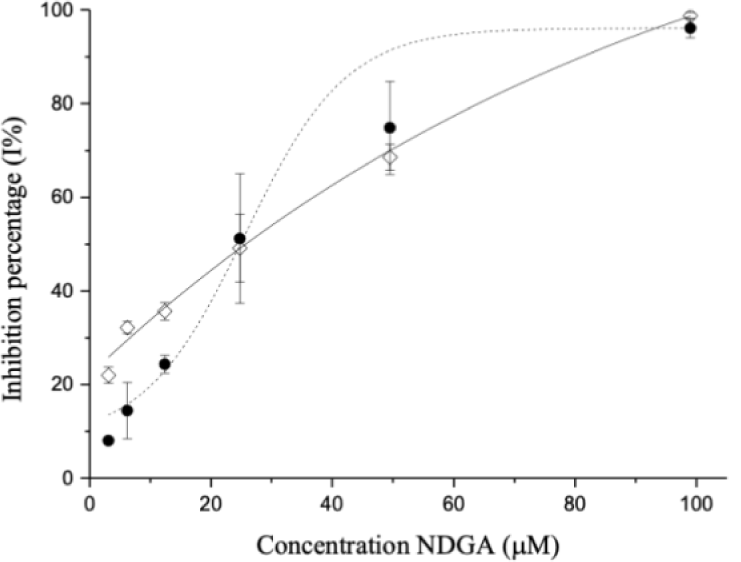
Antiviral activity of NDGA against CbaAr426 (•) and SFCrEq231 (⋄). Inhibition percentage in function of the concentration. The bars represent the 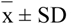 of three independent experiments performed in triplicate.

From these graphics we estimated the EC_50_ (concentration of the treatment able to inhibit 50 % of the viral particles), which was 24.2 ± 1.2 μM and 25.1 ± 6.1 μM for CbaAr426 and SFCrEq231, respectively. The difference between the CE_50_ of both isolates was not significant (*p* = 0.15). Given the CC_50_ in the host cells, it was possible to calculate the selectivity index (SI), which is the relationship between cytotoxicity and activity (SI = CC_50_/EC_50_). Thus, the SI for the NDGA on CbaAr426 was 4.8 and 4.6 for SFCrEq231. It can be observed, in both graphs (Fig. 1) the antiviral activity of NDGA against FSV-like is dose dependent.

### 3.3 Evaluation of the NDGA effect on different times of infection

The significant antiviral activity of NDGA observed against these virus, motivated us to study the NDGA effect at different times of the infection process, produced by the two viral isolates. The initial stages of the infection include the pre-treatment with NDGA during 1 h before the infection (to assess whether the compound has an effect on cells that prevents, in any way, their infection when they are faced with the virus), at the same time of the infection (to evaluate if the compound affects the viral absorption), during the internalization, and 1 h post-internalization (to study if the compound impact in the viral internalization, showing antiviral activity) (Fig. 2). From the results obtained, the %I was calculated for each evaluated condition.

**Figure 2:**
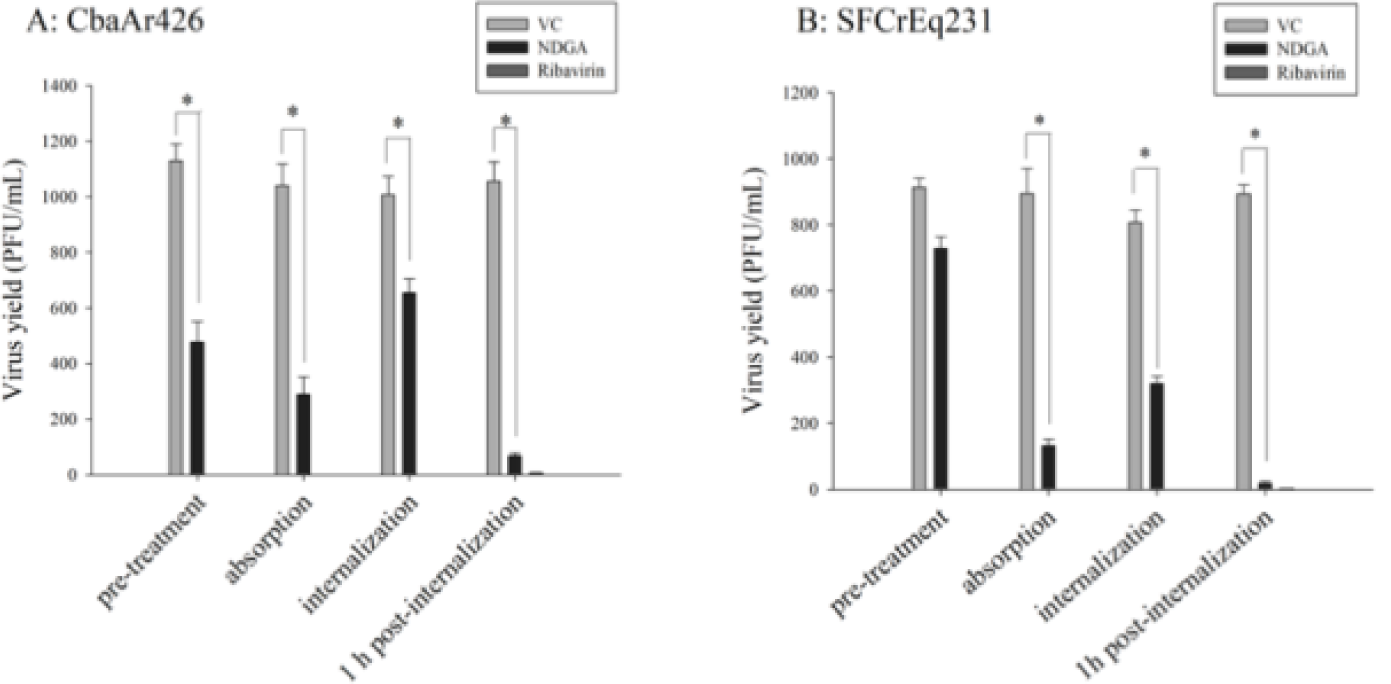
Virus yield (PFU/mL) depending on the treatment with NDGA (MNCC = 90 µM), Ribavirin (90 µM) or without treatment (VC: viral control) at different initial stages of infection with different isolates of FSV-like: CbaAr426 (A) and SFCrEq231 (B). The bars represent the 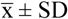 of three independent experiments performed in triplicate. *Significant difference (p < 0.05).

We determined that the infection with CbaAr426 (Fig. 2A) is affected with the NDGA pre-treatment, since 477 ± 74 PFU/mL was observed in comparison with 1130 ± 59 PFU/mL of the VC, which corresponds to 58 %I. Inhibition was also produced during the viral absorption stage (72 I%) and internalization (53 I%), but the most important inhibition was observed when the treatment with NDGA was performed 1 h post-internalization (antiviral activity), showing ≈ 90 I%. On the other hand, in the infection with SFCrEq231 (Fig. 2B), NDGA showed no viral inhibition in the pre-treatment, but it affected the stages of absorption (85 I%) and internalization (60 I%). And also, the greatest inhibition of SFCrEq231 was observed 1 h p.i. (> 95 I%). These results would indicate that the antiviral activity found for NDGA on the isolates of FSV-like is primarily due to an action on the viral particles once they were internalized in the host cell.

To study the effect of NDGA, once the virus was internalized and before a new progeny was released, it was necessary to establish the duration of the virus replication cycle in the used host cells (LLC-MK2). We determined the duration of the virus replication cycle for the two isolates of FSV-like was ≈ 12 h post-infection (supplemental Fig. 4), since extracellular viral particles began to be detected at this time.

To study the effect of the NDGA prior to the release of new virions, we evaluated the NDGA effect every hour during the 8 h p.i., to ensure that the viral particles have not yet exited the host cell. The effect of NDGA was equally important at all tested times (> 90 I%) p.i. of the CbaAr426 isolates (Fig. 3A). However, on the isolates SFCrEq231, the action of the NDGA was only maximal (90 I%) during the first two hours p.i. (Fig. 3B).

**Figure 3:**
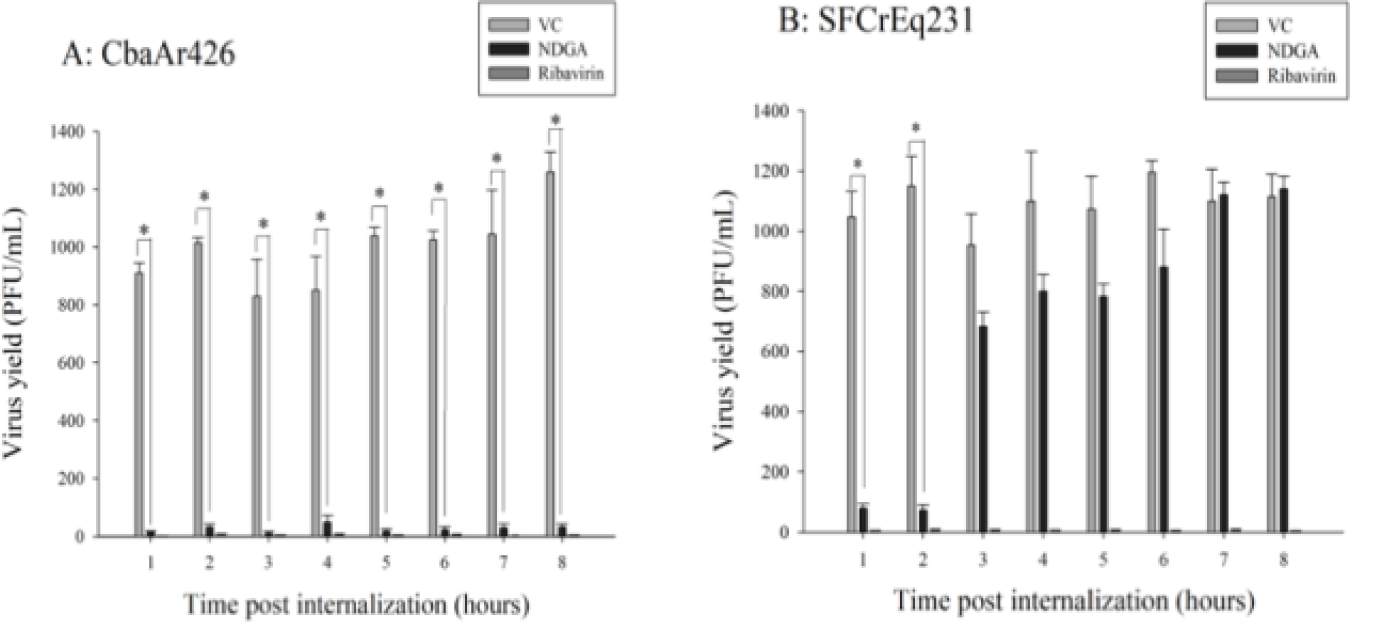
Virus yield (PFU/mL) depending on the treatment with NDGA (MNCC = 90 µM), Ribavirin (90 µM) or without treatment (VC: viral control) at different times post internalization of FSV-like infection with CbaAr426 (A) and SFCrEq231 (B) isolates. The bars represent the 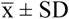 of three independent experiments performed in triplicate. *Significant difference (p < 0.05).

### 3.4 Evaluation of the lipogenesis inhibition in the infection

The antiviral effect of NDGA has been associated with the inhibition of lipogenesis (Syed and Siddiqui, 2011;Soto-Acosta et al., 2014; Merino-Ramos et al., 2017), since NDGA acts as a 5-LOX inhibitor and a negative regulator of the SREBP pathway (Bhattacherjee et al., 1988; Syed and Siddiqui, 2011). To assess the possible involvement of inhibition of the SREBP and 5-LOX pathway in FSV-like infection, specific inhibitors of these two routes were used to establish if their action generates an antiviral effect like NDGA. We selected resveratrol that specifically inhibits SREBP1, the first enzyme in this pathway (Wang et al., 2009), and caffeic acid that is a selective 5-LOX inhibitor (Koshihara et al., 1984).

The cytotoxicity values of resveratrol (Table 2) were obtained from the cell viability curve as a function of concentration (supplemental Fig. 5). In the case of caffeic acid, it showed no cytotoxic effects on LLC-MK2 cells by the neutral red uptake method, being 800 µM the highest concentration tested.

**Table 2:** Values of cytotoxicity for resveratrol and caffeic acid in LLC-MK2 cells by means of the Neutral Red uptake assay.

| | Cytotoxic concentrations (μM) | | |
| --- | --- | --- | --- |
|  | CC<sub>50</sub><sup>1</sup> | SubTC<sup>2</sup> | MNCC<sup>3</sup> |
| Resveratrol | 54.3 ± 3.8 | 18.9 ± 1.4 | 11.6 ± 0.8 |
| Caffeic acid | ND | ND | > 800.000 ± 0.003 |

Data are mean values from three independent experiments $\bar{x} \pm$ SD performed in triplicate.
<sup>1</sup>CC<sub>50</sub>: concentration that reduced the viable cells to 50 %.
<sup>2</sup>SubTC: subtoxic concentration, concentration that reduced the viable cells to 80 % (CC<sub>20</sub>).
<sup>3</sup>MNCC: Maximum Non-Cytotoxic Concentration.
ND: Not detectable

Resveratrol showed no activity on CbaAr426 (Fig. 4B) at the two concentrations tested (SubTC and MNCC). However, it showed a slight inhibition on SFCrEq231 (Fig. 4E), approximately 50 I% at its SubTC. Therefore, we could estimate that the SREBP pathway is involved in the inhibition of the growth of SFCrEq231. However, the observed effect was not as important as that produced by the NDGA (Fig. 4G), where the inhibition was higher than 95 % at its SubTC.

**Figure 4:**
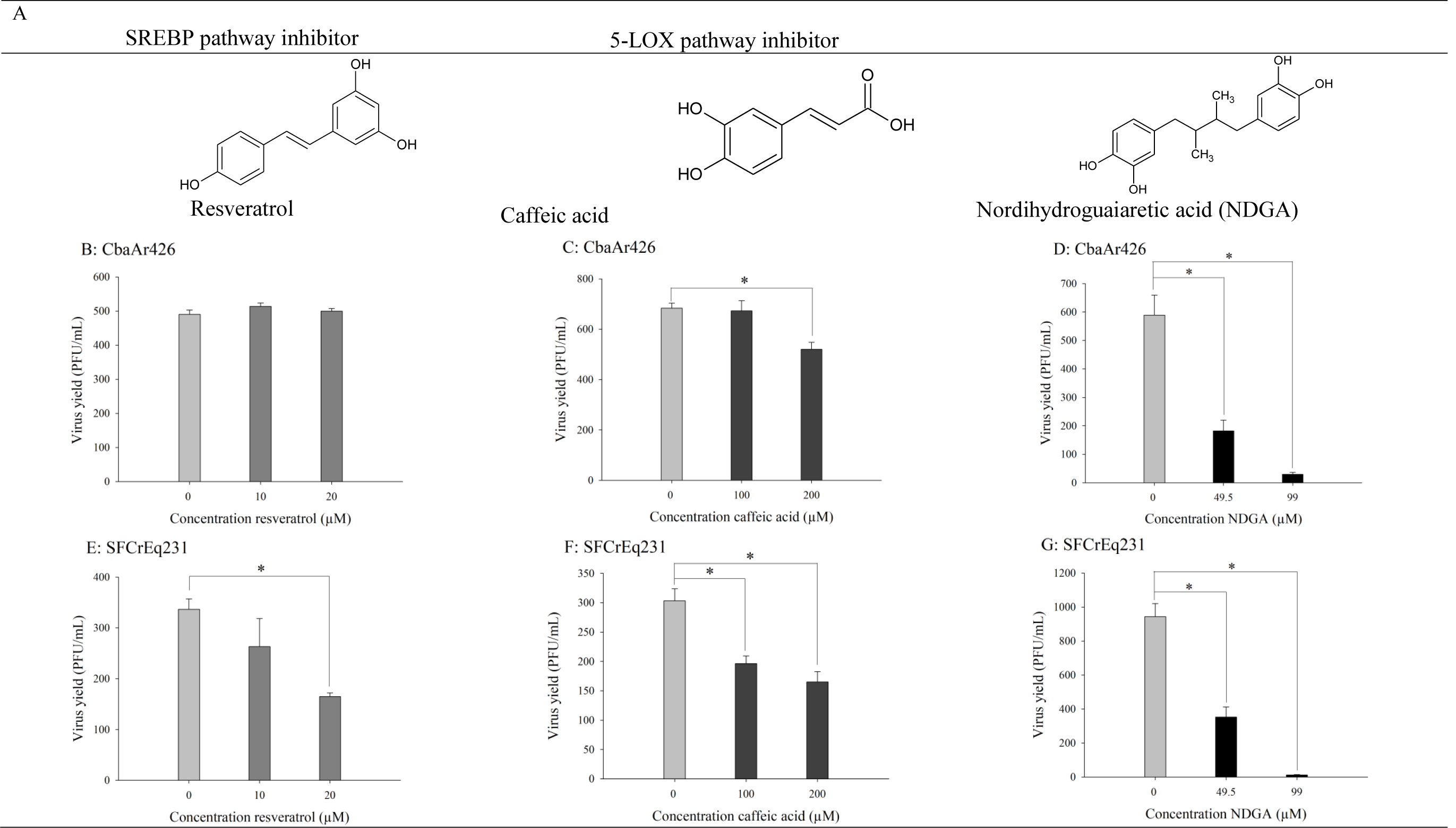
Chemical structure of the different pathway inhibitors and NDGA (A), CbaAr426 yield (PFU/mL) treated with resveratrol (B), caffeic acid (C) and NDGA (D) and yield of SFCrEq231 (PFU/mL) treated with resveratrol (E), caffeic acid (F) and NDGA (G). The bars represent the 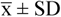 of three independent experiments performed in triplicate. *Significant difference (p < 0.05).

Since caffeic acid was not cytotoxic on host cells, two concentrations were chosen within the non-cytotoxic range (200 and 100 µM) to evaluate the antiviral activity. Caffeic acid showed a mild inhibition on FSV-like isolates at the two tested concentrations (Fig. 4C and 4F), being less than 50 I%. Therefore, we could infer that the 5-LOX pathway would also be involved in the cell cycle of this virus. However, again the inhibitory activity shown by caffeic acid was not as important as that found for NDGA (Fig. 4D and 4G).

### 3.5. NDGA incorporation assay

Having demonstrated that the NDGA does not affect viral particles not associated with cells (virucidal effect) and has the biggest effect after internalization of viral particles, we believe that this compound acts intracellularly. To verify this, we performed an assay to test the ability of NDGA to incorporate into the host cells. The presence and amount of NDGA inside (intracellular fraction) and outside (extracellular fraction) the cells were determined by an HPLC analysis at different times (Table 3).

**Table 3:**
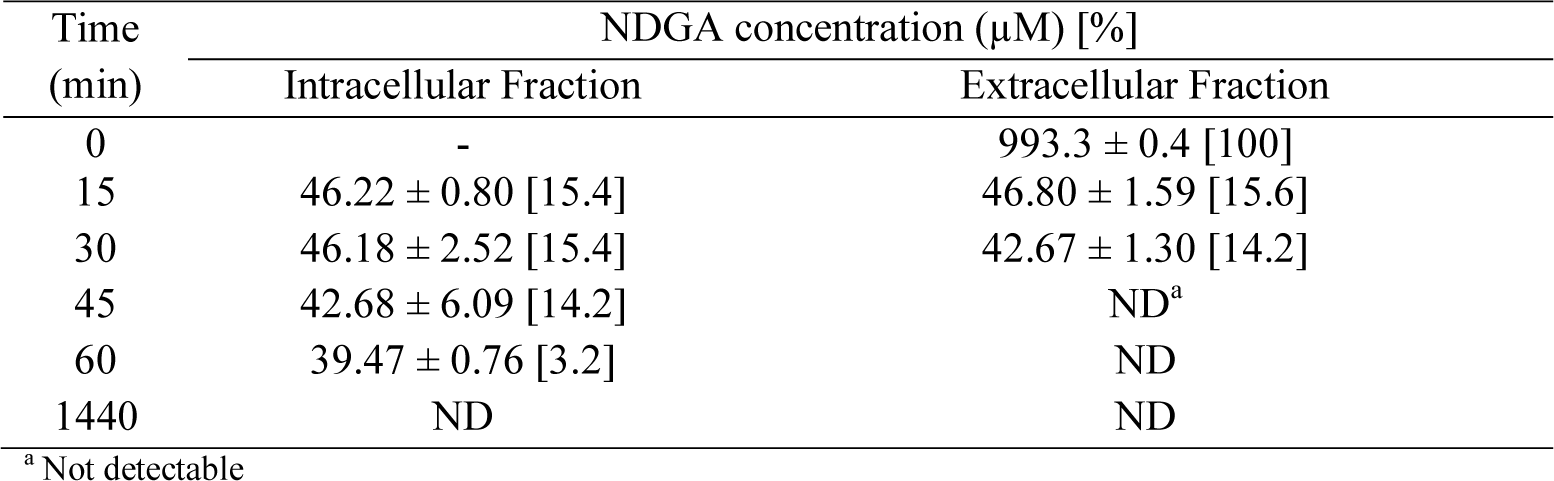
NDGA concentration inside and outside of LLC-MK2 cells at different times.

Results (Table 3) allowed us to verify that NDGA has the ability to enter the host cell (LLC-MK2). The quantification of the NDGA, in both fractions, established that the greatest amount of the compound enters the cell between 15-30 min, being more than 15 % with respect to the amount initially placed. Within the times tested, the presence of NDGA in the intracellular fraction was detected up to 60 min inclusive.

## 4. Discussion

In this work, the *in vitro* antiviral effect of NDGA on two isolates of FSV-like was showed, with a SI value greater than 4. Although there is no a cut-off value for the SI that defines if a natural product is a good antiviral agent, some authors consider that an SI greater than 2 would be sufficient to become a potential antiviral compound (De Clercq, 1993). This result motivated us to deepen the study about the antiviral activity found.

When evaluating the effect of the treatment at initial stages of the FSV-like infection (pre-treatment, absorption and internalization), we established that the inhibitory action of the NDGA was maximum when the treatment was applied 1 h p.i. However, it was remarkable the difference in the behavior of both FSV-like isolates when were treated with NDGA at different times p.i., being the CbaAr426 isolate more susceptible to the NDGA until 8 h p.i. Furthermore, the observed effect of NDGA was comparable to that of ribavirin (p = 0.12).

Given the lack of virucidal activity of NDGA, its greater antiviral activity 1 h p.i. and the verification of its ability to enter the host cell, it can be deduced that the antiviral mechanism of action of this compound against FSV-like would be intracellular.

The antiviral action of NDGA has been associated with an inhibition of lipogenesis that is necessary during the viral replication (Syed and Siddiqui, 2011; Soto-Acosta et al., 2014; Merino-Ramos et al., 2017). Therefore, we study the lipogenesis inhibition by resveratrol (SREBP1 inhibitor) and caffeic acid (5-LOX inhibitor) (Wang et al., 2009; Koshihara et al., 1984) during the infection by FSV-like. Although both inhibitors did not have a reducing effect as high as NDGA, the results indicate that in both FSV-like isolates, the 5-LOX inhibition would be involved in the antiviral effect, whereas the SREBP inhibition was only observed in SFCrEq231 isolate. These results could approximate that FSV-like uses the lipid cell machinery mainly once it is internalized. The relationship between the lipid metabolism of the host cell and the replication of other arbovirus was well characterized (Jordan and Randall, 2016; Osuna-Ramos et al., 2018). Specifically, the antiviral activity of NDGA against HCV and WNV was related to its ability to block the route of the SREBP, which regulates lipid biosynthesis and homeostasis in mammals. NDGA has also shown inhibitory activity on 5-LOX, an enzyme that converts arachidonic acid into oxygenated products with important biological properties (Bhattacherjee et al., 1988). However, there are no reports so far that this interaction occurs in the replication of Orthobunyavirus. Therefore, this work would be a first approximation to possible interactions between this genus replication and cellular lipid metabolism.

The differences in the behavior of both FSV-like isolates when were treated with NDGA at different times and your response to resveratrol, justified the importance of studying two or more different strains; specially when looking for candidate compounds as antivirals, since the same compound could inhibit isolates of the same virus in different ways.

Small molecular differences between strains of the same virus can lead to different behaviors (Nickbakhsh et al., 2019). The CbaAr426 and SFCrEq231 isolates, used in this work, differ in their amino acid sequence, only in Gc glycoprotein encoded in the M segment of the RNA (Tauro et al., 2019). From the three seconds of RNA, the S and L segments of both strains had 100 % homology (GenBank accession nos. **KX100109, KP063894, KX100111**, and **KP063892**). However, there are three differences in the amino acid of the M segment of the two FSV-like isolates, at positions 574/1436 (SFCrEq231: P and CbaAr426: H), 627/1436 (SFCrEq231: I and CbaAr426: T) and 684/1436 (SFCrEq231: N and CbaAr426: S) (GenBank accession nos. **KX100110**, and **KP063896**). These three differences in the amino acid sequence involve the Gc glycoprotein of the FSV-like isolates. So, the differences found between both strains compared to treatment with NDGA and resveratrol may be related to this protein. The possible implication of the Gc glycoprotein from FSV-like in the viral replication cycle or in its relationship with cellular lipid mechanisms, has not been studied in detail. Based on the few available data, it is known that glycoproteins (Gc and Gn) derived from the Orthobunyavirus are responsible for the union and entry of the virus into the cells (Elliott, 2014). M-segment gene products have been implicated in many biological attributes of the virus, including hemagglutination, virulence, tissue tropism, neutralization, and cell fusion (Elliott, 2008). Other authors have already reported that Orthobunyavirus that differ in Gc are associated with different virulence (de Oliveira Filho et al., 2020).

The results of this work lay the foundation for suggesting NDGA as a potential treatment for Orthobunyavirus, where two FSV-like isolates were used as models for the genus. The wide distribution of the Orthobunyavirus, the appearance and reappearance of infections associated with them, and its zoonotic potential highlights the importance of seeking treatment for infection produced by these viruses. It is necessary to emphasize that they are also infections that lack antiviral treatments and vaccine for their prevention.

## Author contributions

F.M. carried out the experimental trials of the entire study, compiled data, drafted the manuscript text with all authors contributing to the drafting process and to the revision of the final manuscript. M.L.M and J.M. They performed the assistance and prepared the equipment for all the analyzes in which it was necessary to use the HPLC. J.J.A. contributed to trials where cells were used, and to the design and supervision of virus experiments, data analysis, and manuscript review. M.S.C., S.C.N.M and B.S.K. She designed and supervised all the work including the manuscript.

## Acknowledgments

We appreciate the contribution of Dr. Guillermo Albrieu Llinas-CONICET, for his contributions in the discussion of this work and Dr. Calos Tonn for providing us the purified nordihydroguaiaretic acid.

The followed funds were used to support the research of this manuscript: FONCYT [BID-PICT 2016 Nº 1697, res. ANPCyT n° 285/17], SeCyT-UNC [2016-2017, res. n° 366/16 y 113/17; Proyecto Consolidar 2018-2021, res. N° 411/18, 99/19; Proyecto Formar, res. N° 411/18, 99/19].

## Appendix A.

Supplementary data

