## Supplementary data for "First report of antiviral activity of nordihydroguaiaretic acid against Fort Sherman-like virus (Orthobunyavirus)"

| | Pages |
|---|---|
| Fig. 1- HPLC-UV chromatogram of NDGA | 2 |
| Fig. 2- Cytotoxicity of ribavirin on LLC-MK2 cells | 3 |
| SA- HPLC analysis | 4 |
| SB- HPLC validation | 5-9 |
| Fig. 3- Cytotoxicity of nordihydroguaiaretic acid in LLC-MK2 cells | 10 |
| Fig. 4- Kinetic curve of replication cycle of two isolates from Forth Sherman-like virus (FSV-like) in the LLC-MK2 cells | 11 |
| Fig. 5- Cytotoxicity of resveratrol on LLC-MK2 cells | 12 |
| SC- References | 13 |

**Supplemental Fig. 1- HPLC-UV chromatogram of NDGA in MeOH**

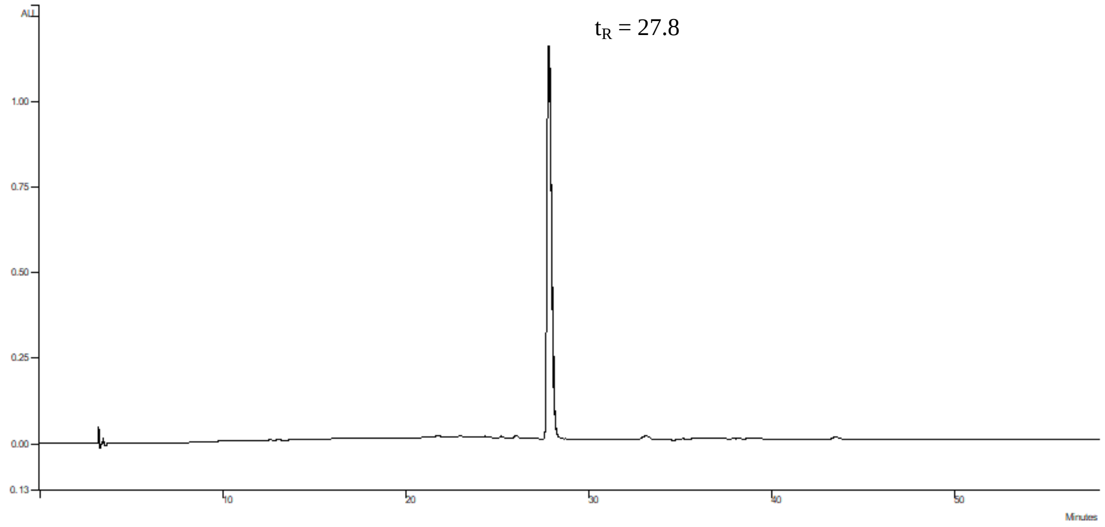

**Supplemental Fig. 2- Cytotoxicity of ribavirin on LLC-MK2 cells.**

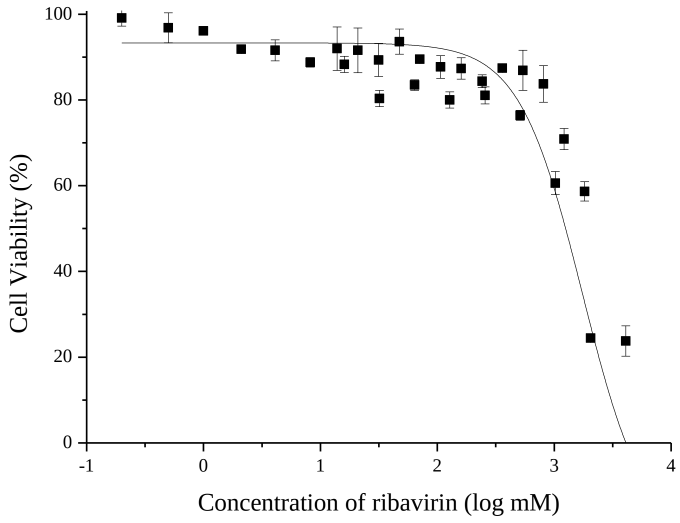


Curves of cellular viability percentage vs. concentration of ribavirin on LLC-MK2 cells, evaluated by NR uptake assay. Each result is expressed as $\overline{x}$ ± SD of at least three independent experiments performed in triplicate. The results were fitted to a sigmoidal dose-response curve, $R^2$ = 0.959.

**Supplemental A- HPLC analysis**

A Varian Pro Star chromatography apparatus (model 210, series 04171, California, USA) equipped with UV-Vis detector was used. The separation was achieved at 25 °C on a Microsorb-MV column 100-5 C-18 (250 x 4.6 mm i.d., Agilent). The mobile phase (1mL/min) involved formic acid in $H_2O$ ultrapure (0.16 M, solvent A) and formic acid in MeOH-HPLC (0.16 M, solvent B), starting with 0% B that changed during 20 min to 70% (10 min), followed by a second ramp (1 min) to 83% B (25 min), returning to the starting conditions for 1 min. Detector was set at 280 nm. The manual injection volume was 20 µL. Data analysis was performed using Varian software (Star Chromatography Workstation 6.41, California, USA). All samples to be analyzed (EC and IC fractions at different times of incubation and standard NDGA) were dissolved in MeOH-HPLC and filtered through paper filter Whatman No. 1 (micro filtration system).

Identification was carried out by comparing the retention times ($t_R$) with the standard NDGA. The $t_R$ values were expressed as means ± SD from three injections for each sample.

Quantitation was achieved by external calibration method, by plotting the area under each peak (AUP) as a function of each NDGA concentration to obtain a calibration curve (7 points). The minimum acceptable value for the correlation coefficient ($R^2$) in the linear regression analysis was 0.99 or more. Standard NDGA was accurately weighed to prepare the corresponding solutions between 3310.9 and 25 µM in MeOH-HPLC, which were submitted to a micro filtration system. Each concentration of NDGA was injected three times, two consecutively and the third a different day for the purpose of method validation.

**Supplemental B- HPLC validation**

Selectivity, linearity, sensibility, and intra-day (repeatability) and inter-day (reproducibility) precision and accuracy of the HPLC method were evaluated, by following the FDA Guidelines (International Conference on Harmonization, 2005a, b). Selectivity implies producing a measurable signal due only to the analyte without interference from other components of the matrix, which was set by choosing a specific wavelength (λ) for NDGA with high absorptivity. Linearity refers to the proportionality between the analyte concentration and its response, which was verified with the compliance of the Lambert-Beer Law on the calibration curve for the range of concentrations tested for the standard NDGA. This parameter was verified with the value of the linear regression coefficient ($R^2 \geq 0.99$) and applying a *t*-student method, where a "*t*" of regression (*tr*) was calculated (Eq. 1), with n-2 degrees of freedom and 95 % of confidence level ($p = 0.025$), whose value must be greater than the tabulated "*t*" value so that the linear correlation is significant in the calculated probability (Quattrocchi *et al*. 1992).

$$tr = \frac{R^2 \cdot \sqrt{(n-2)}}{\sqrt{[1-(R^2)^2]}} \qquad \text{Eq. A.1}$$

Sensitivity corresponds to the smallest amount of analyte that produce a significant result, which was assessed by calculating the Limit of Detection (LOD) and the Limit of Quantitation (LOQ). The values of these limits were determined from the standard deviation (SD) of the Y-axis intersection (Sb) and the slope (a) of the calibration curve achieved by sequentially diluting of a standard solution of NDGA (Quattrocchi *et al*. 1992). It was established that the LOD value must exceed the signal/noise ratio by three SD, and the LOQ must be 10 times this SD.

The intra- and inter-day precision and accuracy of the identification ($t_R$) and quantification (AUP) method were established. From the working curve, two concentrations of NDGA were selected (within the linearity range), which were obtained in triplicate by diluting different stock solutions prepared as described above. These solutions were analyzed in 3 different days (2 of them consecutive), and 3 times every day. The acceptability criteria for intra- and inter-day precision was the confidence interval (n-1, $p = 0.025$), and the relative standard deviation (% RSD) for each tested concentration, whose value should not be higher than the RSD value calculated by the Horwitz equation (LOD was used as concentration in base 10 logarithm) (Horwitz, 1982). The simplest criteria for evaluating intra- and inter-day accuracy are the values obtained for the absolute error (AbsE) and the relative error (RelE), which should be small. A *t*-student method was also performed, where an experimental "*t*" was calculated with n-1 and $p = 0.025$ (Eq. 2). If $texp < t$ of table, the method has the required accuracy for that confidence level, whereas when $texp > t$ of table, the method has a systematic error, of the resulting sign, for that confidence level.

$$texp = \frac{\hat{x}-\bar{x}}{SD\cdot\sqrt{n}} \qquad \text{Eq. A.2}$$

To detect the presence and quantity of NDGA in biological samples, an HPLC method was validated, in which was used a stationary phase and an elution solvent already used for the analysis of extracts of *Larrea* spp (Agüero *et al.*, 2011). Formic acid (0.16 M) was added to the mobile phase to produce a symmetrical signal. To obtain a good resolution in the separation of the eluted compounds, an optimized solvent gradient was developed at the best flow rate.

The selected wavelength for detection (selectivity) corresponds to the maximum absorption of the NDGA (National Library of Medicine), not showing any interference for the compounds in the chromatograms.

The calibration curve performed with the standard NDGA (Fig. 1) complies with the Lambert-Beer Law (linearity) in the range of tested concentrations (25 to 3310.9 μM), with a value of $R^2 > 0.999$ and a calculated value of *tr* (49.9) greater than the tabulated value (2.571, n-2, $p = 0.025$).

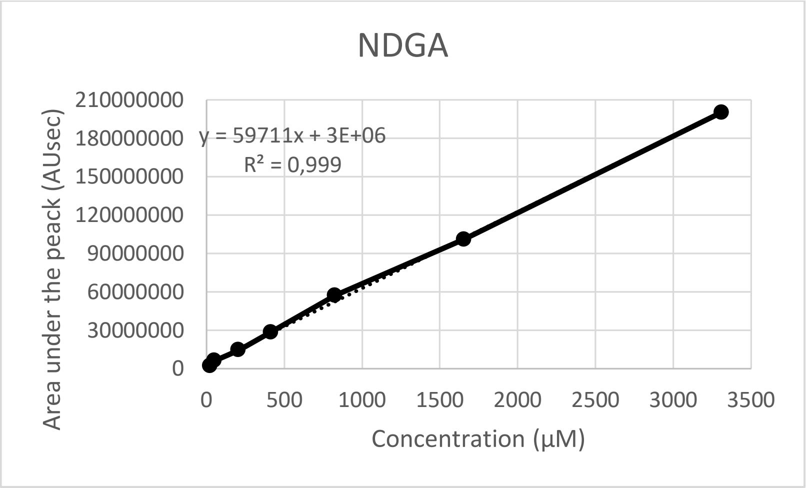


Fig. B1: Calibration curve of NDGA

The parameters that allow evaluating the sensitivity of the method were calculated, setting the Limit of detection (LOD) in 8.32 μM (lowest concentration of analyte that can be detected in a sample, but not necessarily can be quantified) and the Limit of quantification (LOQ) in 11.3 μM ( lowest analyte concentration that can be determined with a reasonable precision and accuracy).

The intra- and inter-day precision of the identification ($t_R$) and quantification (AUP) method were established by the values of the relative standard deviation (% RSD) of two solutions of NDGA (200 and 25 μM), which were analyzed in 3 different days (2 of them consecutive), and 3 times every day. The results (Table 1 and 2) showed a good repeatability and reproducibility in the method for the identification and quantification of NDGA, since the the intra- and inter-day % RSD were lesser than the maximum RSD calculated by the Horwitz equation.

Table B1: Intra- and inter-day precision for the identification of NDGA

| Solution | Intra-day precision (Repeatability) | | | | | | % RSD by Horwitz | Inter-day precision (Reproducibility) | |
|---|---|---|---|---|---|---|---|---|---|
| | Day 1 | | Day 2 | | Day 3 | | | | |
| | $t_R$ (min) | % RSD | $t_R$ (min) | % RSD | $t_R$ (min) | % RSD | | $t_R$ (min) | % RSD |
| **1** | 27.57 ± 0.02[a] | 0.07 | 27.6 ± 0.1[a] | 0.36 | 27.7 ± 0.1[a] | 0.36 | 1.45 | 27.6 ± 0.1[d] | 0.36 |
| **2** | 27.7 ± 0.1[a] | 0.36 | 27.82 ± 0.01[a] | 0.04 | 27.6 ± 0.2[a] | 0.72 | | 27.7 ± 0.1[d] | 0.36 |
| **1** and **2** | 27.6 ± 0.1[b] | 0.36 | 27.7 ± 0.1[b] | 0.36 | 27.6 ± 0.1[b] | 0.36 | | 27.7 ± 0.1[e] | 0.36 |

a) n = 3, b) n = 6, c) LOD was used as concentration in base 10 logarithm, d) n = 9, e) n = 18

Table B2: Intra- and inter-day precision for the quantification of NDGA

| Solution | **Intra-day precision** (Repeatability) | | | | | | % RSD by Horwitz | **Inter-day precision** (Reproducibility) | |
|---|---|---|---|---|---|---|---|---|---|
| | Day 1 | | Day 2 | | Day 3 | | | | |
| | µM | % RSD | µM | % RSD | µM | % RSD | | µM | % RSD |
| **1** | 202 ± 2[a] | 1 | 201 ± 2[a] | 1 | 202 ± 1[a] | 0.5 | 1.45[b] | 202 ± 2[c] | 1 |
| **2** | 24.8 ± 0.2[a] | 0.8 | 25.5 ± 0.1[a] | 0.4 | 25.0 ± 0.2[a] | 0.8 | | 25.1 ± 0.3[c] | 1.2 |

a) n = 3, b) LOD was used as concentration in base 10 logarithm, c) n = 9,

The intra- and inter-day accuracy of the identification ($t_R$) and quantification (AUP) method were determined by the small values obtained for the absolute error (AbsE) and the relative error (RelE), with the resulting sign, for a 95 % confidence level. In addition, a *t*-student method was also carried out, in which if the value of the experimental "*t*" (*texp*, n-1, $p = 0.025$) was lesser than the value of tabulated "*t*" means that the method has the required accuracy for the confidence level used (95 %). The results showed (Table 3 and Table 4) that the method has the required accuracy for the identification and quantification of NDGA.

Table B.3: Intra- and inter day accuracy for the identification of NDGA

| **Intra-day accuracy** (Repeatability) (n = 3) | | | | | **Inter-day accuracy** (Reproducibility) (n = 9) | | | |
|---|---|---|---|---|---|---|---|---|
| Day | AbsE | RelE | *texp* | *t* | AbsE | RelE | *texp* | *t* |
| 1 | 0.15 | 0.54 | 0.24 | 4.303 | -0.01 | 0.04 | 0.007 | 2.306 |
| 2 | -0.15 | 0.55 | 0.11 | | | | | |
| 3 | -0.03 | 0.12 | 0.13 | | | | | |

AbsE: absolute error, RelE: relative error, *texp*: value of *t* experimental, *t*: tabulated *t* (n = 1, $p = 0.025$)

Table B.4: Intra- and inter day accuracy for the quantification of NDGA

| **Solution** | **Intra-day accuracy** (Repeatability) (n = 3) | | | | | **Inter-day accuracy** (Reproducibility) (n = 9) | | | |
|---|---|---|---|---|---|---|---|---|---|
| | Day | AbsE | RelE | *texp* | *t* | AbsE | RelE | *texp* | *t* |
| **1** | 1 | 0.17 | 0.1 | 0.05 | 4.303 | -0.46 | 0.3 | 0.07 | 2.306 |
| | 2 | -1.34 | 0.7 | 0.34 | | | | | |
| | 3 | -0.22 | 0.1 | 0.05 | | | | | |
| **2** | 1 | -0.06 | 0.2 | 0.11 | | -0.04 | 0.15 | 0.04 | |
| | 2 | -0.22 | 0.9 | 0.45 | | | | | |
| | 3 | 0.17 | 0.7 | 0.35 | | | | | |

AbsE: absolute error, RelE: relative error, *texp*: value of *t* experimental, *t*: tabulated *t* (n = 1, *p* = 0.025)

In conclusion, we consider that this HPLC method is suitable for the detection and quantitation of NDGA in biological samples.

**Supplemental Fig. 3- Cytotoxicity of nordihydroguaiaretic acid in LLC-MK2 cells**

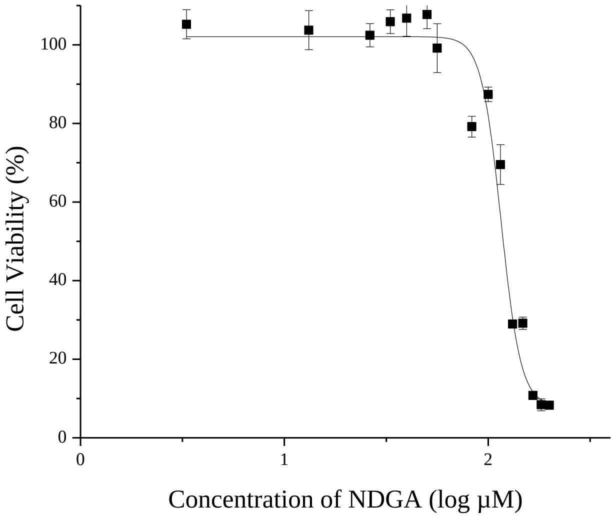


Curves of cellular viability percentage vs. concentration of NDGA on LLC-MK2 cells, evaluated by NR uptake assay. Each result is expressed as $\overline{x} \pm SD$ of at least three independent experiments performed in triplicate. The results were fitted to a sigmoidal dose-response curve, $R^2 = 0.968$.

**Supplemental Fig. 4- Kinetic curve of replication cycle of two isolates from Forth Sherman-like virus (FSV-like) in the LLC-MK2 cells**

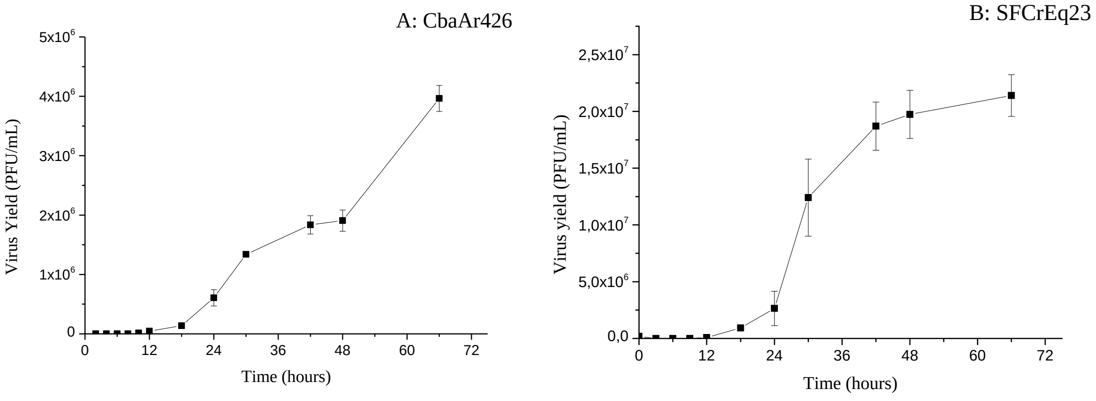


Kinetics of replication of CbaAr426 (A) and SFCrEq231 (B) in LLC-MK2 by Plate-forming units (PFU/mL) of the extracellular supernatants. The bars represent the $\bar{x} \pm$ SD of two independent experiments performed in triplicate.

**Supplemental Fig. 5- Cytotoxicity of resveratrol on LLC-MK2 cells**

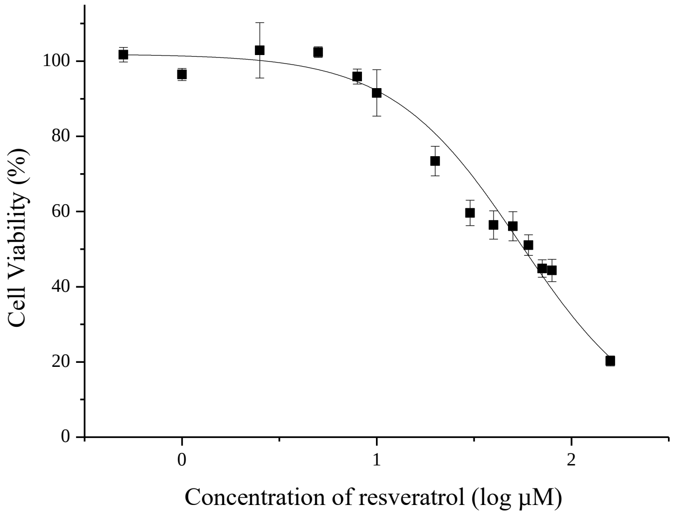


Curves of cellular viability percentage vs. concentration of resveratrol on LLC-MK2 cells, evaluated by NR uptake assay. Each result is expressed as $\bar{x} \pm SD$ of at least three independent experiments performed in triplicate. The results were fitted to a sigmoidal dose-response curve, $R^2 = 0.981$.

**Supplemental C- References**

National Library of Medicine (NIH). Nordihydroguiaretic acid. Available in: https://pubchem.ncbi.nlm.nih.gov/compound/Nordihydroguaiaretic-acid [29th june, 2020].

Agüero, M.B., Svetaz, L., Sánchez, M., Luna, L., Lima, B., López, M.L., Zacchino, S., Palermo, J., Wunderlin, D., Feresin, G.E., Tapia, A., 2011. Argentinean Andean propolis associated with the medicinal plant *Larrea nitida* Cav. (*Zygophyllaceae*). HPLC–MS and GC–MS characterization and antifungal activity. *Food Chem. Toxicol*. 49, 1970–1978.
